# Cryo-EM structures of Arabidopsis PRC2 histone methyltransferase isoforms reveal a differential regulatory mechanism

**DOI:** 10.64898/2026.06.22.733897

**Authors:** Kyeong-Seon Hong, Junghyun Kim, Sibum Sung, Ji-Joon Song

## Abstract

Polycomb Repressive Complex 2 (PRC2) is a histone H3K27 methyltransferase that represses gene expression. Arabidopsis thaliana (A. thaliana) has several different PRC2 isoforms that are functionally distinct during the life cycle of the plants. However, their biochemical and structural characteristics have not been investigated. Here, we biochemically characterized PRC2 isoforms having different catalytic subunits: SWNINGER (SWN; PRC2_SWN_) and CURLY LEAF (CLF; PRC2_CLF_). Interestingly, PRC2_SWN_ showed much lower activity than PRC2_CLF_. In addition, PRC2_SWN_ methylates histone H3K27 in mono and di-methylation, while PRC2_CLF_ shows robust tri-methylase activity. We also determined the cryo-electron microscopy (cryo-EM) structures of PRC2_SWN_ and PRC2_CLF_, revealing that the substrate binding pocket of the SWN SET domain is blocked by a loop in the pre-SET domain, functioning as an auto-inhibitory loop, while that of the CLF SET domain is freely accessible. Introduction of CLF-like mutations in the auto-inhibitory loop in SWN enhances PRC2_SWN_ activity. Furthermore, structure-guided in planta analysis shows that a CLF-mimetic SWN mutant rescues the CLF knockout phenotype. Our work provides structural and molecular insights into the isoform-specific regulatory mechanism of plant PRC2.

## Introduction

In eukaryotes, DNA forms into high-order structure called chromatin with a fundamental unit of nucleosome composed of 146 bp DNA and a histone octamer (Luger et al. 1997). Various epigenetic factors alter the chromatin structure, including chromatin remodeling, histone chaperoning, and histone post-translational modifications (PTMs) (Allis and Jenuwein 2016). Among these, histone PTMs play crucial roles in the epigenetic regulation of gene expression (Strahl and Allis 2000). The PTMs alter chromatin states in a manner dependent on the chemical moiety (e.g., methylation, acetylation, phosphorylation, and ubiquitination) and the positions of modifications (Kouzarides 2007). Among these, histone H3 lysine 27 methylation (H3K27), exclusively catalyzed by PRC2, acts as a key repressive mark in developmental processes (Margueron and Reinberg 2011). PRC2 is highly conserved across eukaryotes, from *Drosophila* to humans. In humans, it comprises a core complex consisting of EZH2 or EZH1 (the catalytic subunit containing the SET domain), EED, RBAP46/48, and scaffold subunit SUZ12 (Cao et al. 2002; Muller et al. 2002; Margueron and Reinberg 2011). Similarly, plant PRC2, composed of a highly conserved four-subunit core complex, functions as a master epigenetic regulator governing key life-cycle transitions, including vegetative growth, flowering, vernalization, and seed development (Mozgova and Hennig 2015). Consequently, loss-of-function mutations in essential PRC2 subunits lead to severe developmental defects (Goodrich et al. 1997; Yoshida et al. 2001; Chanvivattana et al. 2004). Interestingly, plant PRC2 exhibits a more complex assembly, utilizing varied combinations of three catalytic subunit homologs (SWN, CLF, and MEA in *Arabidopsis*) and three scaffold subunit homologs (EMF2, VRN2, and FIS2 in *Arabidopsis*), together with the core component FIE and MSI (Derkacheva and Hennig 2014). Each catalytic subunit seems to have its distinct role in mediating H3K27 methylation. For instance, while SWN is associated with broader genome-wide spreading of the mark, it has only minor effects on global methylation levels and developmental phenotypes. Conversely, CLF exhibits more restricted spreading but exerts a profound impact on global methylation and plant phenotype (Baerenfaller et al. 2011; Shu et al. 2019). Moreover, SWN and CLF are distinctly involved in the biphasic establishment of H3K27me3 in *Arabidopsis* - nucleation and spreading (Costa and Dean 2019). Both SWN and CLF participate in the initial nucleation, which occurs at loci proximal to the transcription start site (TSS). However, during the subsequent spreading phase to more distal sites from TSS following cold exposure, CLF acts exclusively as the sole methyltransferase (Yang et al. 2017). Despite that plant PRC2 variants containing distinct catalytic subunits play differential roles in regulating plant developmental processes, our understanding of their biochemical and structural characteristics remains limited. Here, we biochemically dissected *Arabidopsis* PRC2 variants with the SWN or CLF catalytic subunits, revealing their distinct methyltransferase activities. Furthermore, our cryo-EM structures of PRC2_SWN_ and PRC2_CLF_ revealed an auto-inhibitory mechanism in SWN, wherein a loop within the pre-SET domain blocks the SET domain in SWN but not CLF. The functional importance of this auto-inhibitory loop was subsequently validated through *in vitro* and *in planta* studies. Collectively, our findings provide structural and molecular insights into the variant-specific mechanisms underlying the function of plant PRC2 isoforms.

## Results

### Biochemical characterization of Arabidopsis PRC2 isoforms

To characterize the biochemical activities of plant PRC2 variants, we successfully purified two full-length PRC2 isoforms having different catalytic subunits (SWN and CLF): PRC2_SWN_ (composed of FIE, SWN, EMF2 and MSI) and PRC2_CLF_ (composed of FIE, CLF, EMF2, MSI) (Fig. 1A). We then measured the histone methyltransferase (HMTase) activities of the PRC2 complexes using G5E4 nucleosomal arrays (Fig. 1B). Interestingly, PRC2_CLF_ showed much higher activity than PRC2_SWN_. To systematically characterize the PRC2 activities, we performed kinetic analysis (Fig. 1C). Both PRC2_CLF_ and PRC2_SWN_ show similar K_m_ values (0.27±0.07 μM for PRC2_CLF_ and 0.37±0.07 μM for PRC2_SWN_), but PRC2_CLF_ (Vmax 35.1±8.4) has higher V_max_ value than PRC2_SWN_ (Vmax 16.5±3.2), indicating that the catalytic properties intrinsically differ between PRC2_CLF_ and PRC2_SWN_. We then examined the methylation status of histone H3K27 by these complexes (Fig.1D). PRC2_SWN_ methylates histone H3K27 up to di-methyl while PRC2_CLF_ exhibited mono-, di- and tri-methylation activity. Human PRC2 are allosterically activated by trimethylated H3K27 (Margueron et al. 2009; Poepsel et al. 2018). Therefore, we also tested whether the tri-methylated histone H3K27me3 peptide stimulates the plant PRC2. While PRC2_SWN_ showed clear allosteric activation effect by addition of H3K27me3 peptide, PRC2 _CLF_ exhibited marginal activation effect (Supplemental Fig. S1A). These data imply that the plant PRC2 isoform has distinct catalytic property and the catalytic subunits (SWN and CLF) govern this property.

**Figure 1.**
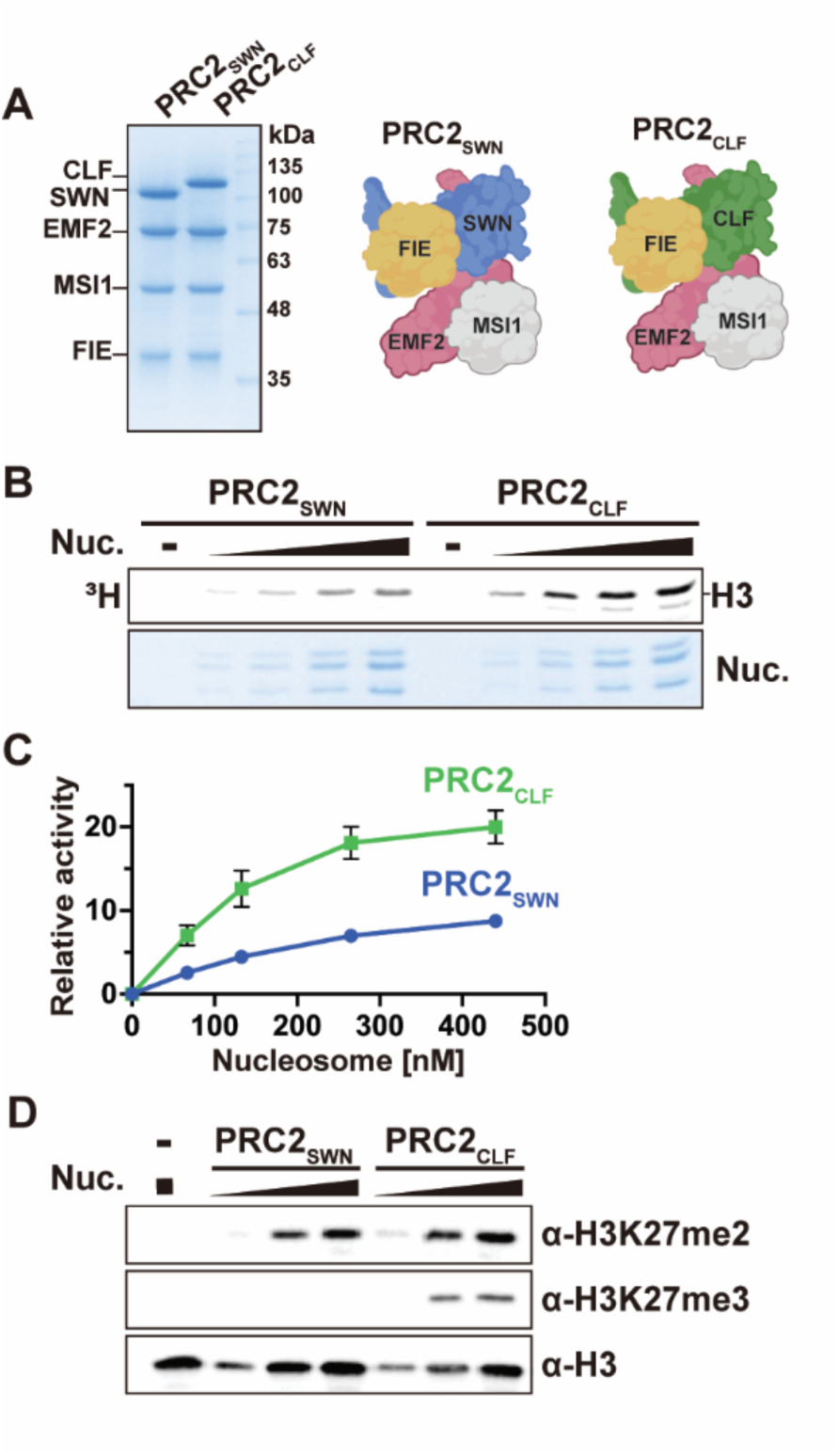
Biochemical characterization of Arabidopsis PRC2 variants. (*A*) An SDS-PAGE gel of the purified PRC2_SWN_ and PRC2_CLF_ and schematic illustrations representing PRC2 subunits of each variant. (*B*) Histone methyltransferase assay (HMTase assay) of PRC2_SWN_ and PRC2_CLF_ with G5E4 nucleosomal arrays (Nuc). Autoradiographs for methylated histone H3 Lys27 with ^3^H-SAM (top panel) and a Coomassie-stained membrane (bottom panel) used for auto-radiography signal detection showing histones. (*C*) Kinetic analysis of PRC2 HMTase activity. *V_max_* (PRC2_SWN_: 16.5±3.2, PRC2_CLF_: 35.1±8.4) and *K_m_* (PRC2_SWN_: 0.37±0.07 μM, PRC2_CLF_: 0.27±0.07 μM) values of each variant were calculated using Lineweaver-Burk plot. (*D*) Western-blot analysis of H3, H3K27me2 and H3K27me3 levels following an *in vitro* HMTase assay with PRC2_SWN_ and PRC2_CLF_.

### Cryo-EM structures of PRC2_SWN_ and PRC2_CLF_

To understand the molecular mechanism for the differential activities of the PRC2 isoforms, we determined the cryo-EM structures of PRC2_SWN_ and PRC2_CLF_. As we failed to determine the structures of the full-length PRC2, we decided to use a truncated version of EMF2 subunit. Human PRC2 with SUZ12-VEFS domain is sufficient to form a PRC2 complex without RbAp48 and showed comparable activity to the full-length PRC2 complex (Jiao and Liu 2015; Justin et al. 2016). Therefore, we generated ΔPRC2_SWN_ (EMF2_505-631 a.a._) and ΔPRC2_CLF_ (EMF2_495-631 a.a._) without MSI subunits (Supplemental Fig. S2A). We then validated that the EMF2 truncated PRC2 have a similar activity to the full-length PRC2 (Supplemental Fig. S2B).

We collected a total of 12,506 and 13,516 micrographs for ΔPRC2_SWN_ and ΔPRC2_CLF_ respectively using a Titan Krios 300 keV microscope with a Falcon4i direct detector. We then processed the data and determined the cryo-EM structures of ΔPRC2_SWN_ and ΔPRC2_CLF_ in 3.67 Å and 3.5 Å resolutions respectively (Fig. 2; Fig. 5; Supplemental Figs. S3; Supplemental Figs. S4; Supplemental Table S1). The model building of PRC2 complexes was guided by AlphaFold (Jumper et al. 2021). The overall structures of ΔPRC2_SWN_ and ΔPRC2_CLF_ are similar to each other and also to that of human PRC2 (Supplemental Fig. S6). In our cryo-EM structures, the VEFS domain of EMF2 acts as a scaffold linking the catalytic subunits (SWN and CLF) and FIE subunit. The FIE subunit interacts with the EMF2-VEFS domain and the N-terminal SANT Binding Domain (SBD), EED binding domain (EBD) and β-addition motif (BAM) domains of the catalytic subunits (CLF, SWN). The SANT1 domain in ΔPRC2_SWN_ is well structured while it is missing in the ΔPRC2_CLF_ cryo-EM map. We modeled in the cofactor, SAM for visualization purpose. Our cryo-EM structures show the highly conserved structures between plant and human PRC2.

**Figure 2.**
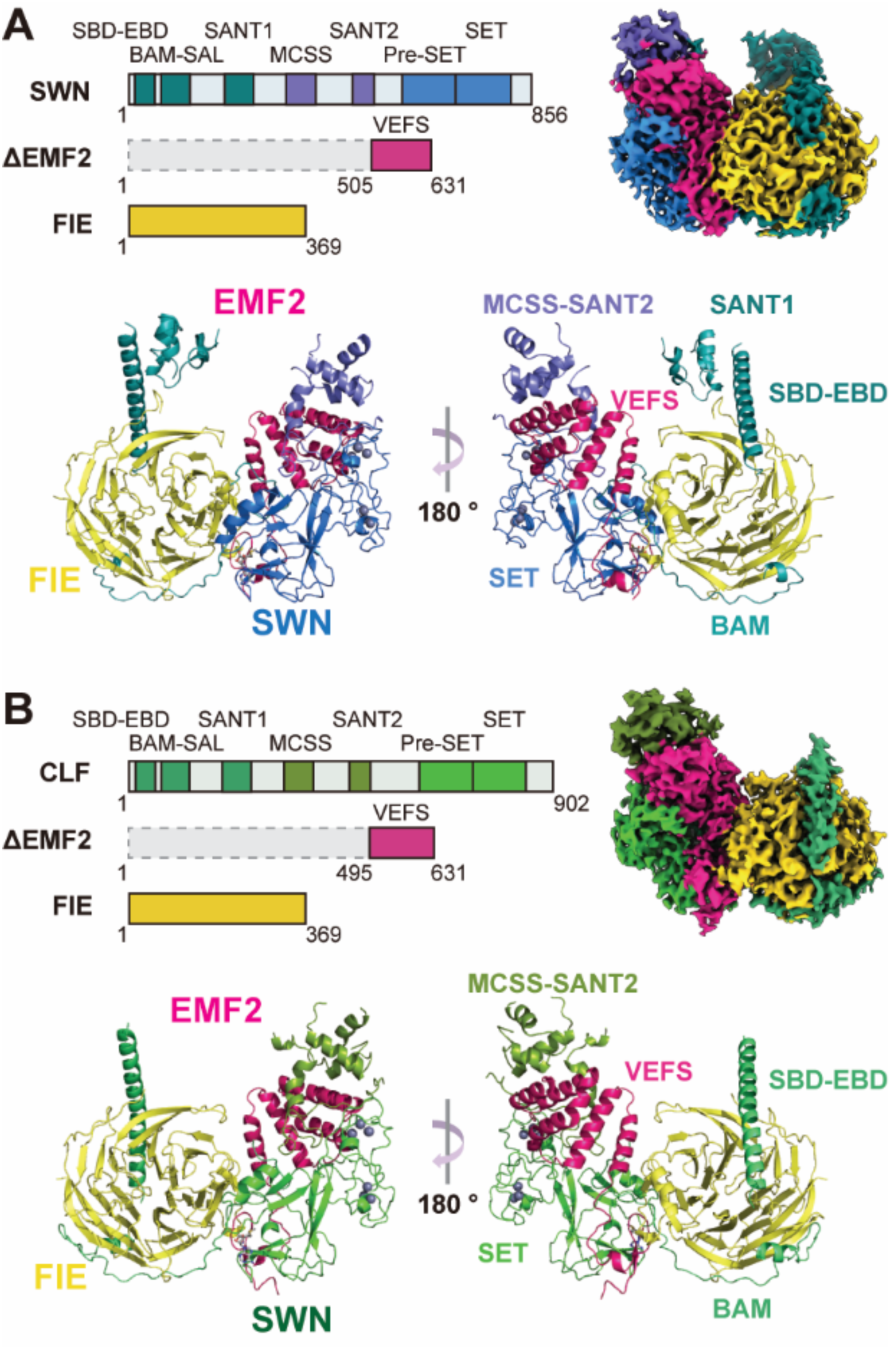
Cryo-EM structures of PRC2_SWN_ and PRC2_CLF_. (*A*) (*B*) Schematic diagrams of the constructed used for PRC2_SWN_ (*A*) and PRC2_CLF_ (*B*) structures (top left panels). The cryo-EM maps of PRC2_SWN_ and PRC2_CLF_ (top right panels of *A, B*). The atomic structures of the PRC2_SWN_ (*A*) and PRC2_SWN_ (*B*), are shown in ribbon diagram (bottom panels of *A, B*). EMF2 is colored magenta, FIE in yellow, and individual domains of the SWN and CLF are colored according to the color shown in the schematic diagram (*A*) and (*B*).

### The auto-inhibitory loop in the pre-SET domain blocks the substrate binding pocket of SWN but not CLF

Despite the similarity between the PRC2_SWN_ and PRC2_CLF_ structures, there is a striking difference in the SET catalytic domain. In the SET domain of SWN, but not CLF, a loop from the pre-SET domain occupies the substrate binding pocket, as if the loop functions as an auto-inhibitory (AI) loop (Fig. 3A). Although the auto-inhibitory mechanism by the post-SET auto-inhibitory loop was reported in isolated SET domains including EZH2, NSD2 and ASH1L (Qiao et al. 2011; Antonysamy et al. 2013; Wu et al. 2013; Rogawski et al. 2015; Lee et al. 2019), to our knowledge, this is the first observation of the auto-inhibitory loop from the pre-SET domain. While the catalytic pocket is blocked by the AI loop in the SWN, the corresponding loop in CLF is disordered. Therefore, the substrate binding pocket of CLF is freely accessible (Figs. 3A). These observations are consistent with that PRC2_CLF_ showed higher activity than PRC2_SWN_ (Fig. 1). The AI loop (_592_SVWKRIAG_599_) in the SWN pre-SET domain is well ordered as clearly visible in the cryo-EM (Supplemental Fig. S5). The AI loop interacts with the rest of the SET domain of SWN (Fig. 3B). Most interestingly, K595 in the AI loop is inserted into the active site where the histone H3K27 normally occupies. The hydrocarbon region of K595 in the AI loop interacts with Y821 as the H3K27 of histone H3 does. In addition, W594 interacts with a hydrophobic patch formed by L761 and V769 from the β5- and β6-strands, and A746 form the SET-I helix. In addition, R596 in the AI loop interacts with D763 in the SET domain. Although there is no sequence similarity between histone H3 and the AI loop except the lysine, the AI loop occupies the substate binding pocket blocking the access of the substrate (Fig. 3C and Fig. 3D). These data suggest that the AI loop regulates PRC2_SWN_ activity.

**Figure 3.**
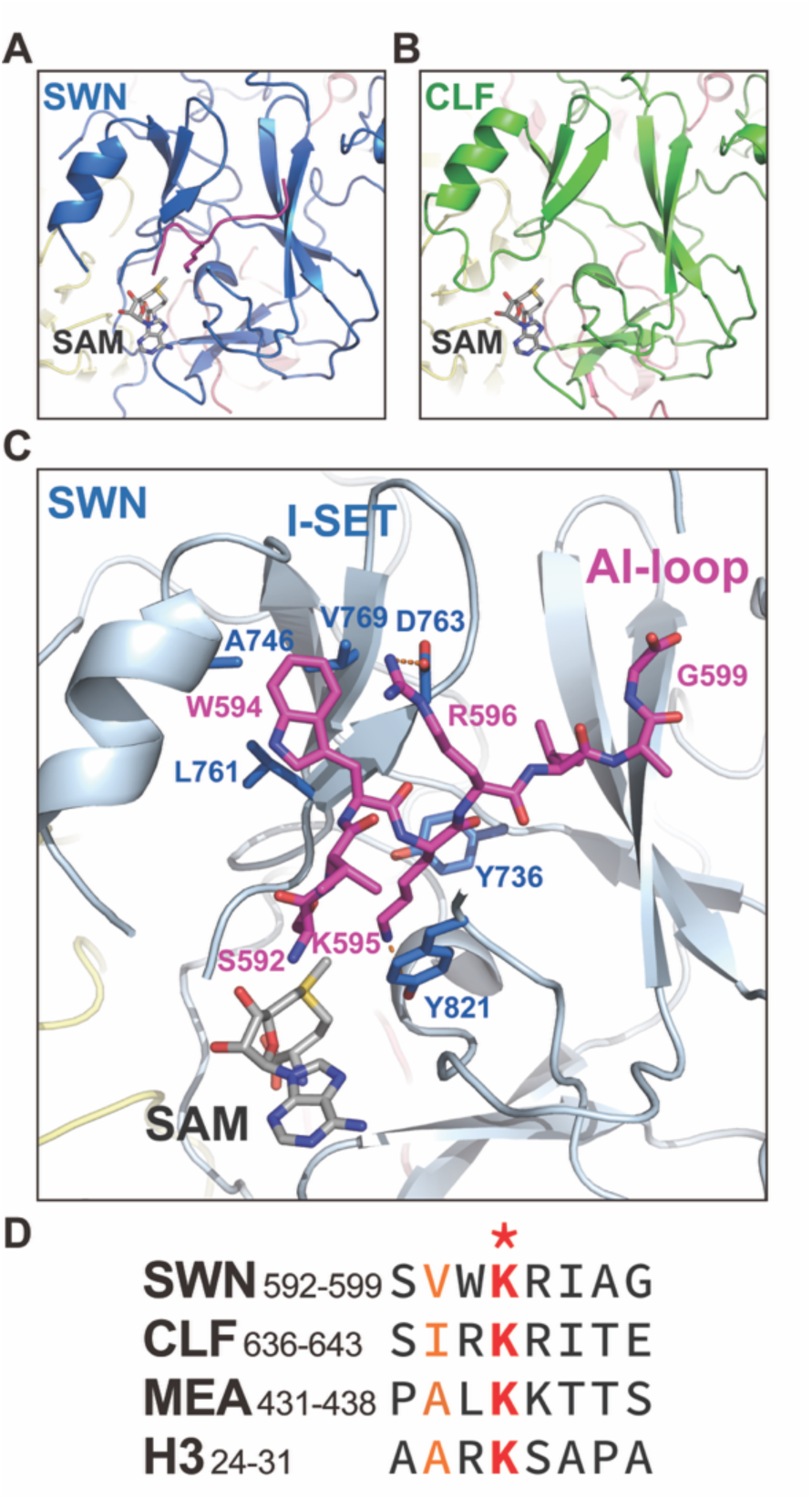
The substrate binding pocket of SWN, but not CLF, is blocked by an auto-inhibitory (AI) loop in the Pre-SET domain. (*A*) The AI loop (shown in magenta color) occupies the histone H3 binding pocket in the SWN SET domain. (*B*) The substrate binding pocket of CLF SET is freely accessible. S-Adenosyl-Methionine (SAM)s were modeled in the cryo-EM structures for visualization of the substrate binding pocket based on the human PRC2 structure (PDB: 7TD5) for *A, B*. (*C*) The detailed interaction between the AI loop and SWN SET domain. W594 of the AI-loop interacts with the hydrophobic pockets formed by A746, L761, V769 in the SWN SET domain. R596 of the AI-loop forms a salt bridge with D763 in the SET domain. Lys594 of the AI-loop is inserted into the catalytic pocket and interacts with Y821 of the SET domain. (*D*) Sequence alignment of the AI-loop region among SWN, CLF, MEA and H3. The lysine residues (SWN K595, CLF K639, MEA K434 and H3 K27) are highlighted in red with asterisk mark. Other conserved residues in these loops are highlighted in orange.

### Swapping of the auto-inhibitory loops between SWN and CLF enhances PRC2_SWN_ activity

The AI loop is marginally conserved between SWN and CLF (Fig 3D). Although the KRI motif in the AI loop exists exist both in SWN and CLF, the flanking sequences of the KRI motif differ. To examine the role of the AI loop, we generated full-length chimera SWN and CLF having the AI loops swapped (Fig. 4A). We replaced the AI loop of SWN with the corresponding loop of CLF to generate full-length PRC2_SWN_AI-CLF_ and conversely replaced the corresponding loop of CLF with the AI loop of SWN to generate full-length PRC2_CLF_AI-SWN_ (Supplemental Fig. S7). We then performed HMTase assays using these chimeric PRC2 complexes (Fig. 4B). Interestingly, the PRC2_SWN_AI-CLF_ exhibited substantially higher HMTase activity than PRC2_SWN_, indicating that the inhibitory effect of the AI loop was released in PRC2_SWN_AI-CLF_. However, PRC2_CLF_AI-SWN_ showed comparable activity to that of the PRC2_CLF_, suggesting that the AI loop is not sufficient to block the substrate binding pocket of CLF. We further examined the methylation status catalyzed by the chimeric PRC2s (Fig. 4C). Although PRC2_SWN_AI-CLF_ displayed significantly enhanced activity relative to PRC2_SWN_, this enhancement was restricted to mono- and di-methylation, with no increase in tri-methylation. In contrast, PRC2_CLF_AI-SWN_ showed similar overall HMTase activity to PRC2_CLF_, with predominant di- and tri-methylation. These results suggest that AI loop swapping enhances overall PRC2_SWN_ activity while preserving its intrinsic catalytic preference for mono- and di-methylation. Overall, these data further support that the AI loop in SWN regulates the activity.

**Figure 4.**
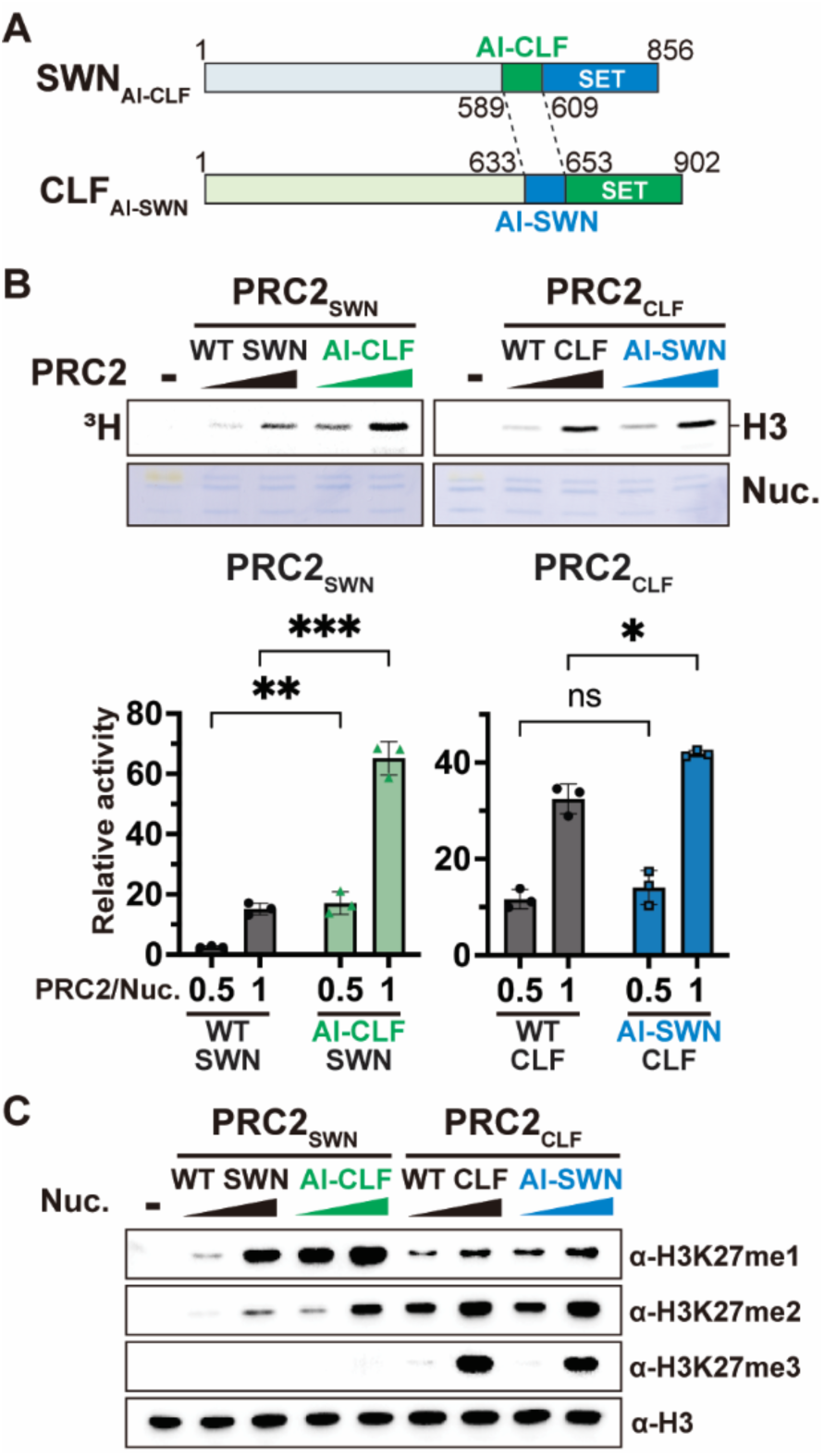
A loop swapping between SWN and CLF simulates SWN HMTase activity. (*A*) A schematic diagram of the AI-loop swapped SWN and CLF. The AI loop (589-609 a.a.) of SWN and the corresponding loop (633-653 a.a.) of CLF were swapped with each other. (*B*) HMTase assay with wild-type PRC2 (WT SWN, WT CLF) and the loop swapped PRC2 (PRC2_SWN_/AI-CLF and PRC2_CLF_/AI-SWN). Autoradiographs for methylated histone H3K27 with ^3^H-SAM (top panel) and a Coomassie-stained membrane (bottom panel) used for auto-radiography signal detection. Bar graphs with quantification of relative activity of HMTase assay (bottom panel). In the bar graphs, each dot represents a biological replicate (*n*=3), error bars represent the mean ± SD. *P*-values were calculated with multiple t-test and adjusted using False Discovery Rate (FDR) method (*: *p* < 0.05, **: *p* < 0.01, ***: *p* < 0.001). (*C*) Western-blot analysis of H3, H3K27me1, H3K27me2 and H3K27me3 levels on the product from HMTase assay with PRC2s using G5E4 nucleosomal array.

### SWN_AI-CLF sufficiently complements the clf phenotype in Arabidopsis

To determine whether PRC2_SWN_AI-CLF_ can complement the *clf* phenotype, we generated transgenic *Arabidopsis thaliana* lines expressing either *CLF*, *SWN*, and *SWN_AI-CLF* in the *clf* mutant background and evaluated their phenotypes (Figs. 5A and Supplemental Fig. S8A). The *clf* mutant phenotype is well characterized, exhibiting dwarfism, early flowering, and small, curled leaves in *Arabidopsis*, mainly due to misregulation of developmental regulators such as *AGAMOUS (AG)* (Goodrich et al. 1997). Since primary transgenic plants exhibit considerable phenotypic variation, we measured rosette diameter immediately after bolting as a quantitative parameter to distinguish *clf*-like plants from phenotypically rescued plants. Approximately 15% of the primary transgenic lines expressing wild-type CLF rescued the *clf* phenotype (Fig. 5A and Supplemental Fig. S8A). Interestingly, 7.7% of the primary transgenic lines expressing SWN_AI-CLF also exhibited rescue of the *clf* phenotype, whereas none of the lines expressing wild-type SWN showed phenotypic rescue (Fig. 5A; Supplemental Fig. S8A). This difference was not due to variation in transgene expression or protein enrichment. Both SWN and SWN_AI-CLF accumulated to comparable levels of the enrichment at the target locus, indicating that loop swapping did not alter chromatin association of the protein (Supplemental Fig. S8B).

**Figure 5.**
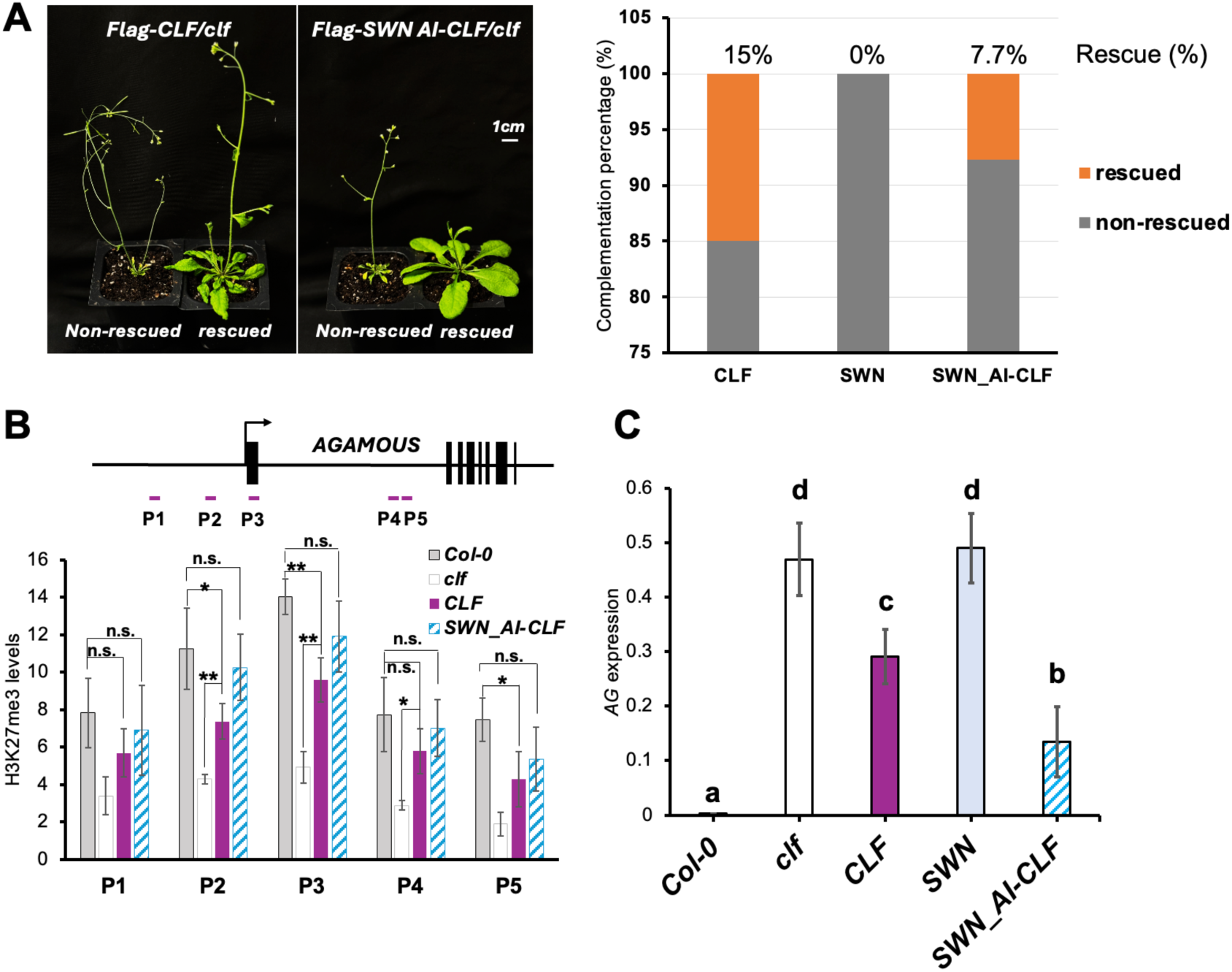
Transgenic plants expressing SWN_AI-CLF fully complement the *clf* phenotype in Arabidopsis. (*A*) Phenotypic complementation of the *clf* mutant. Representative images of T1 transgenic lines expressing either *Flag-CLF* or *Flag-SWN_AI-CLF* in the *clf* mutant background. Morphological phenotypes are categorized as non-rescued (*clf* like) or rescued. Scale bar = 1 cm. Bar graph showing the complementation percentage of transgenic groups exhibiting non-rescued (*clf* like) or fully rescued phenotypes across three different independent transgenic lines. (*B*) A schematic representation of ChIP-qPCR amplicons at the *AG* locus and ChIP-qPCR analysis comparing H3K27me3 levels at the *AG* locus across indicated genotypes. The H3K27me3 levels were normalized against the level of *rDNA* internal control. Error bars indicate ± SD from three biological replicates. P-values were calculated using a two-tailed Student’s t-test (n.s.: not significant, *: *p* <0.05, **: *p* <0.01). n.s. indicates not significant. (*C*) Relative expression levels of *AG*. Transcript levels of *AG* were normalized against the internal control *PP2A* and represent the mean ± SD from three biological replicates. Different letters above the bars indicate statistically significant differences determined by one-way ANOVA followed by Tukey’s honestly significant difference (HSD) post-hoc test (P<0.05).

To further assess functional rescue, we examined H3K27me3 levels at the *AG* locus, a shared target of CLF and SWN (Schubert et al. 2006; Shu et al. 2019) in the rescued transgenic lines expressing *CLF* and *SWN_AI-CLF*, using chromatin immunoprecipitation (ChIP) assays with H3K27me3 antibodies (Kuo and Allis 1999) (Fig. 5B). Consistent with the phenotypic rescue observed in the CLF and SWN_AI-CLF lines, H3K27me3 levels at *AG* were restored to near wild-type levels in both transgenic backgrounds. Correspondingly, *AG* expression was significantly repressed in the CLF and SWN_AI-CLF lines relative to the *clf* mutant (Fig. 5C). Together, these results demonstrate that SWN_AI-CLF acquires CLF-like function in vivo and support a model in which SWN activity is regulated by its auto-inhibitory loop.

## Discussion

CLF and SWN exhibit substantial functional redundancy. However, genetic studies have demonstrated that they also possess distinct biological functions. Loss of CLF results in severe developmental defects, including early flowering and ectopic expression of floral homeotic genes, whereas *swn* single mutants display relatively mild phenotypes (Shu et al. 2019). Nevertheless, SWN can partially compensate for CLF function at many target loci, and only the *clf swn* double mutant exhibits a nearly complete loss of H3K27me3 and severe developmental abnormalities (Shu et al. 2019). These observations suggest that CLF and SWN share overlapping functions but also possess intrinsic properties that confer locus-specific regulatory activities. However, the molecular basis underlying these functional differences remains poorly understood. In this study, we delineated the structural and molecular characteristics of plant PRC2 isoforms having different catalytic subunits (SWN and CLF).

Our data showed the biochemical distinction between SWN and CLF closely parallels that observed between EZH1 and EZH2. PRC2_CLF_ efficiently catalyzes mono-, di-, and tri-methylation of H3K27 as PRC2_EZH2_. In contrast, PRC2_SWN_ resembles PRC2_EZH1_, exhibiting substantially weaker catalytic activity, predominantly generating mono- and di-methylation (Margueron et al. 2008; Lee et al. 2018). It is also notable that PRC2_SWN_ and PRC2_EZH1_ maintains H3K27me3 at the nucleation site while PRC2_CLF_ and PRC2_EZH2_ are required for the spreading of H3K27me3 (Yang et al. 2017). However, PRC2_SWN_ and PRC2_EZH2_ are significantly stimulated by its product, H3K27me3 but PRC2_CLF_ and PRC2_EZH1_ exhibited a marginal stimulation. These observations suggest that plant PRC2 isoforms have distinct properties to human PRC2.

The auto-inhibitory mechanism was observed in other SET domains. However, in all case, the AI loop from the post-SET domain, not pre-SET domain, occupies in the catalytic pocket in isolated SET domain (An et al. 2011; Qiao et al. 2011; Antonysamy et al. 2013; Wu et al. 2013). These AI loops are released upon the complex formation (Jiao and Liu 2015; Hou et al. 2019; Lee et al. 2019). However, in plant SWN SET domain, the AI loop exists in the pre-SET domain and the auto-inhibitory state is maintained in the context of the complex. However, this auto-inhibitory state was not observed in the CLF SET domain. These observations suggests that the different catalytic subunit isoforms in the plant PRC2 have intrinsically distinct regulatory mechanism.

Interestingly, the AI loop in SWN interact with the SET-I helix, which was shown to mediate the allosteric activation by binding of histone H3K27me3 on EED via the stimulation-responsive motif (SRM) in human PRC2 (Poepsel et al. 2018). Consistent with this, we also observed the allosteric activation of PRC2_SWN_ by histone H3K27me3 peptide, but not in PRC2_CLF._ This allosteric effect may facilitate the release of the AI loop. In addition, the K595 in the AI loop mimicking the H3K27 is inserted in the catalytic pocket, raising a possibility that the K595 can be also methylated. The modification of the AI loop may regulate to release the AI loop.

While the overall activity PRC2_SWN_AI-CLF_ exhibited higher activity than PRC2_SWN_, the stimulated activity is limited to mono- and di-methyltransferase similar to PRC2_SWN_ in-vitro. However, SWN_AI-CLF fully rescued the *clf* phenotype, suggesting that SWN_AI-CLF can generate tri-methylated histone H3K27 in-vivo. At this moment, there is no clear explanation on how SWN_AI-CLF fully methylation in-vivo. There are several proteins identified to interact with plant PRC2 to regulate its activity including VIVIPAROUS1/2 (VAL1/2), VERNALIZATION INSENSITIVE 3 (VIN3), VERNALIZATION INSENSITIVE 3-Like 1 (VIL1)/VRNALIZATION 5 (VRN5), LIKE HETEROCHROMATIN PROTEIN 1 (LHP1) and others (Sung and Amasino 2004; Veluchamy et al. 2016; Zhou et al. 2017; Godwin and Farrona 2022; Franco-Echevarria et al. 2023). In addition, a recent study showed that ULTRAPETALA1 stimulate PRC2_SWN_ but not PRC2_CLF_ activities (Geshkovski et al. 2026).

Therefore, it is plausible that binding of PRC2 interacting proteins may alter the PRC2 activities to activate the activity by releasing of the AI loop. However, at this moment, it is unclear how the AI loop is released from the catalytic site and PRC2_SWN_ get activated, and further structural and biochemical study will be necessary to understand the exact mechanism by which the auto-inhibition is relieased. In summary, our works provide structural and mechanistic insights on the regulatory mechanism of plant PRC2 activity also imply that plant PRC2 isoforms have complicate and delicate regulatory mechanism, which to be further discovered.

## Materials and Methods

### Expression and purification of Arabidopsis PRC2 complexes

The components of *Arabidopsis thaliana* PRC2: SWN (residues 1–856), CLF (residues 1–902), EMF2 (full-length, residues 1–631), truncated EMF2 variants (VEFS domain; residues 495–631 or 505–631), MSI1 (residues 1–424), and FLAG-tagged FIE (residues 1–369) were cloned into pFastBac1 vectors. Point mutations were generated using the KOD Plus Mutagenesis Kit (Toyobo, Japan). Bacmids were generated by transformation into DH10Bac cells and purified for baculovirus production. Recombinant baculoviruses were subsequently generated and amplified to passage 3 in *Spodoptera frugiperda* (Sf9) cells. For protein expression, passage 3 baculoviruses encoding individual PRC2 subunits were co-infected into Sf9 cells at a density of 3.5–4.0 × 10^6^ cells/mL. Cells were harvested 48 h post-infection and resuspended in lysis buffer containing 100 mM NaCl, 50 mM Tris-HCl (pH 8.0), 5% glycerol, and cOmplete ULTRA protease inhibitor cocktail (Roche, Switzerland). Cells were lysed by three freeze–thaw cycles, and cellular debris was removed by centrifugation at 27,216 × g for 2 h. The clarified supernatant was incubated with anti-DYKDDDDK G1 affinity resin (GenScript, USA) for 2 h at 4°C. The resin was subsequently washed with five column volumes of wash buffer containing 100–500 mM NaCl, 50 mM Tris-HCl (pH 8.0), and 5% glycerol. Bound proteins were eluted using elution buffer containing 0.3 mg/mL FLAG peptide, 100 mM NaCl, 50 mM Tris-HCl (pH 8.0), and 5% glycerol. For cryo-EM sample preparation, the N-terminal FLAG tag on FIE was removed by overnight incubation with TEV protease at 4°C at a ratio of 0.2 mg TEV protease per 4 mg of target protein in buffer containing 100 mM NaCl, 50 mM Tris-HCl (pH 8.0), 0.5 mM EDTA, and 1 mM DTT. The protein samples were further purified by ion-exchange chromatography using a HiTrap Q HP column (Cytiva, USA) with a linear NaCl gradient from 50 mM to 1 M, followed by size-exclusion chromatography on a Superdex 200 Increase 10/300 GL column (Cytiva, USA) equilibrated in buffer containing 100 mM NaCl, 50 mM Tris-HCl (pH 8.0), and 5% glycerol. Fractions containing the PRC2 complex were pooled and concentrated using Amicon Ultra centrifugal filters with a 100 kDa molecular weight cutoff (MWCO) (Merck Millipore, USA) to a final concentration of 1.0–2.0 mg/mL. The purified protein samples were flash-frozen in liquid nitrogen and stored at −80°C until use.

### Histone methyltransferase assay (HMTase assay)

Recombinant *Xenopus laevis* histones H2A, H2B, H3, and H4 were expressed, purified, and lyophilized as previously described (Luger et al. 1999). Briefly, lyophilized histones were individually dissolved in unfolding buffer containing 6 M guanidine hydrochloride (GnHCl), 20 mM Tris-HCl (pH 7.5), 0.25 mM EDTA, and 5 mM DTT, and subsequently mixed at a molar ratio of H2A:H2B:H3:H4 = 1.1:1.1:1:1. Histone octamers were assembled by dialysis against refolding buffer containing 2 M NaCl, 10 mM Tris-HCl (pH 7.5), 1 mM EDTA, and 5 mM β-mercaptoethanol at 4°C overnight. The assembled histone octamers were further purified by size-exclusion chromatography using a HiLoad 26/600 Superdex 200 pg column (Cytiva, USA). For nucleosome array assembly, purified histone octamers were mixed with G5E4 DNA for 12-mer nucleosome array reconstitution at a 1:1 mass ratio and assembled by stepwise salt dialysis (Owen-Hughes and Workman 1996; Ikeda et al. 1999).

For histone methyltransferase (HMTase) assays, PRC2 variants (1.4 μM) were incubated with G5E4 nucleosome arrays at the indicated PRC2:nucleosome molar ratios in reaction buffer containing 66 mM NaCl, 50 mM Tris-HCl (pH 8.0), 4 mM DTT, and 13.5 μM [^3^H]-S-adenosyl-L-methionine ([^3^H]-SAM; Revvity, USA) for 1.5 h at 25°C. For H3K27me3 peptide stimulation assays, PRC2 complexes at a seven-fold lower concentration (0.2 μM) were incubated with 0.1 μM G5E4 nucleosome arrays and H3K27me3 peptides at molar ratios ranging from 1:5 to 1:20 relative to PRC2 under identical reaction conditions. For assays using auto-inhibitory loop mutants, 0.1–0.2 μM PRC2 was incubated with 0.1 μM G5E4 nucleosome arrays under the same conditions. Reactions were terminated by addition of 4 μL of 5× SDS sample buffer followed by heating at 95°C for 10 min. Samples were separated on 12% SDS-PAGE gels and subsequently transferred onto Immobilon-PSQ membranes (Millipore, USA) by semi-dry transfer in buffer containing 25 mM Tris, 190 mM glycine, 20% methanol, and 0.01% SDS. Following air-drying, membranes were exposed to phosphor imaging plates, and radioactive signals were detected using an FLA-7000 Fuji BAS imager (Fujifilm, Japan). The intensities of the ^3H signals were quantified using Image Lab software.

### Western blot

For Western blot analysis, the reaction and electrophoresis conditions were identical to those used for the HMTase assays described above, except that non-radioactive (cold) S-adenosyl-L-methionine (SAM; Sigma-Aldrich, USA) was used in place of [^3^H]-SAM. Following SDS-PAGE, proteins were transferred onto Immobilon-PSQ membranes (Millipore, USA) using a wet-transfer system in transfer buffer containing 25 mM Tris, 190 mM glycine, 10% methanol, and 0.01% SDS. The methylation status of histone H3 was analyzed using antibodies against H3K27me1 (Abcam, ab194688), H3K27me2 (Abcam, ab24684), H3K27me3 (Abcam, ab192985), and total histone H3 (Abcam, ab1791). Immunoreactive bands were detected using an HRP-conjugated anti-rabbit IgG secondary antibody (Cell Signaling Technology, 1:3,000 dilution) and visualized using Western Blot Chemiluminescent Substrates (Bionics).

### Cryo-EM grid preparation

The ΔPRC2_SWN_ and ΔPRC2_CLF_ complexes were cross-linked with 0.5 mM bis(sulfosuccinimidyl) suberate (BS3; Thermo Fisher Scientific, USA) for 25 min at 25°C in buffer containing 100 mM NaCl, 25 mM Tris-HCl (pH 8.0), and 1 mM DTT. The cross-linked complexes were further purified by size-exclusion chromatography using a Superdex 200 Increase 10/300 GL column (Cytiva, USA) equilibrated with buffer containing 100 mM NaCl and 50 mM Tris-HCl (pH 8.0). Fractions containing the PRC2 complexes were pooled and concentrated using Amicon Ultra centrifugal filters with a 100 kDa molecular weight cutoff (Merck Millipore, USA) in the presence of 0.03% *n*-octyl-β-D-glucoside (β-OG; Sigma-Aldrich, USA) to a final protein concentration of approximately 0.2 mg/mL. For cryo-EM grid preparation, 3 μL of each sample was applied to Quantifoil R1.2/1.3 Cu 300 mesh holey carbon grids coated with graphene oxide (0.085 mg/mL, prepared in-house). Grids were plunge-frozen using a Vitrobot Mark IV (Thermo Fisher Scientific) operated at 4°C and 100% relative humidity with a blotting time of 3 s and a blot force of −12.

### Data collection and processing

Cryo-EM data were collected using a Titan Krios G4 300 keV transmission electron microscope (Thermo Fisher Scientific, USA) equipped with a Falcon 4i direct electron detector (Thermo Fisher Scientific, USA) at the KAIST Analysis Center for Research Advancement (KARA). For the determination of the ΔPRC2_SWN_ structure, 13,516 movie stacks were collected at a nominal magnification of 105,000x, corresponding to a calibrated pixel size of 0.97 Å. Images were acquired with a total electron dose of 50 e^-1^/Å^2^ and a defocus range of −1.2 to −2.8 μm. Data processing was performed using CryoSPARC v4.3.1 (Punjani et al. 2017). Movie frames were corrected using Patch Motion Correction, which is based on MotionCor2 (Zheng et al. 2017), and contrast transfer function (CTF) parameters were estimated using Patch CTF, which incorporates Gctf (Zhang 2016). After quality assessment based on CTF fit resolution, defocus, astigmatism, and full-frame motion, 9,102 micrographs were selected for further processing. Template-based particle picking using a low-resolution preliminary PRC2 map identified 2,129,509 particles. Subsequent two-dimensional (2D) classification reduced the dataset to 1,064,802 particles. Initial three-dimensional (3D) reconstructions were generated by *ab initio* reconstruction, and the best class containing 557,290 particles was selected for further refinement. Three rounds of non-uniform refinement combined with local motion correction (Rubinstein and Brubaker 2015) and global and local CTF refinement, yielded a final reconstruction at 3.67 Å resolution based on the gold-standard Fourier shell correlation (FSC) criterion of 0.143, with an estimated B-factor of 212.8 Å^2^. For the determination of the ΔPRC2_CLF_ structure, 12,506 movie stacks were collected under identical imaging conditions, including a nominal magnification of 105,000x (pixel size of 0.97 Å), a total dose of 50 e^-1^/Å^2^, and a defocus range of −1.2 to −2.8 μm. Following motion correction and CTF estimation, 9,176 micrographs were selected for further analysis. Template-based particle picking yielded 1,650,760 particles, which were reduced to 938,011 particles after 2D classification. The selected ab initio reconstruction containing 406,693 particles was subsequently refined through iterative rounds of non-uniform refinement and CTF refinement. The final reconstruction reached a global resolution of 3.30 Å according to the FSC = 0.143 criterion, with an estimated B-factor of 166.6 Å^2^.

### Model building

Initial atomic models of each PRC2 variant were generated using AlphaFold predictions (Jumper et al. 2021). The predicted models were rigid-body fitted into the corresponding cryo-EM density maps using ChimeraX v1.8 (Pettersen et al. 2021). The fitted models were subsequently manually adjusted and refined to optimize map-to-model agreement and stereochemical quality using Phenix (Liebschner et al. 2019), Coot (Emsley et al. 2010), and ISOLDE implemented in ChimeraX (Croll 2018). Model validation was performed by assessing stereochemical parameters, including Ramachandran statistics, rotamer outliers, Cβ deviations, and atomic clashes. Detailed refinement and validation statistics are summarized in Supplementary Table S1. The cryo-EM maps and corresponding atomic coordinates have been deposited in the Electron Microscopy Data Bank (EMDB) and the Protein Data Bank (PDB). The ΔPRC2_SWN_ structure has been deposited under EMDB accession code EMD-81511 and PDB accession code 27WJ, whereas the ΔPRC2_CLF_ structure has been deposited under EMDB accession code EMD-81512 and PDB accession code 27WK.

### Plant Materials and Growth Condition

*Arabidopsis thaliana* plants utilized in this study were of the Columbia-0 (Col-0) background and cultivated under a continuous temperature of 22℃ in a growth room with a 16h light/8h dark long-day (LD) cycle. For each transgenic line, the coding DNA sequences (CDS) of *CLF*, *SWN*, and the loop-swapped *SWN* variant (SWN_AI-CLF) was cloned into the pGWB612 vector (containing the 35S promoter and an N-terminal Flag tag). *clf* (*clf-29*) mutant plants were transformed via the floral dip method using *Agrobacterium tumefaciens* (strain GV3101) cultures harboring each construct (T0), and subsequent transformants were selected on Basta-containing medium (T1). A total of 65∼72 independent T1 transgenic lines for each construct were selected for phenotypic analysis. Rosette diameters were measured immediately upon bolting.

### Chromatin Immunoprecipitation (ChIP) assay

10-day-old seedlings grown at 22℃ under long-day conditions (16h light/8h dark) were collected and crosslinked in 1% formaldehyde under vacuum for 15 min (5 min on - break - 5 min on – break – 5 min on), then quenched by adding glycine to a final concentration of 0.125 M under vacuum for 5 min. Crosslinked tissue was rinsed three times with distilled water, excess liquid was removed, and samples were flash-frozen and ground in liquid nitrogen. Nuclei were isolated and the chromatin was sonicated using a Bioruptor at high power to obtain ∼200–500 bp DNA fragments. For immunoprecipitation, chromatin was incubated with anti-H3K27me3 antibody (Millipore, 07-449) followed by capture with Dynabeads Protein A (Invitrogen), while anti-FLAG M2 affinity gel (Sigma-Aldrich, A2220) was used for FLAG-tagged samples. After extensive washing, protein–DNA complexes were eluted, crosslinks were reversed, and DNA was purified. Enrichment of target regions was quantified by quantitative real-time PCR (qPCR) using gene-specific primers (Supplemental Table S2). ChIP signals were normalized to input DNA and expressed as fold enrichment relative to a control region (*rDNA*). Three biological replicates were performed for H3K27me3 and SWN ChIP-qPCR assays.

### RNA analysis

10-day-old seedlings grown at 22℃ under long-day conditions (16h light/8h dark) were collected and total RNA was extracted utilizing the TRIzol (Invitrogen) method. The extracted RNA was subsequently treated with DNase (Promega) at 37℃ for 30 minutes to eliminate residual genomic DNA before reverse transcription. For complementary DNA (cDNA) synthesis, 2 µg of total RNA was hybridized with oligo(dT) primers at 65℃ for 5 min. Reverse transcription was then performed using M-MLV reverse transcriptase (Invitrogen) at 37℃ for 50 min, followed by enzyme inactivation at 70℃ for 15 min.

Quantitative real-time PCR (qPCR), employing specific primer sets (Supplemental Table S2), was utilized to determine the transcript levels of target genes, with normalization to *PP2A*.

## Acknowledgement

We thank the member of the Song laboratory for helpful discussion. This work is supported by grants from National Research Foundation of Korea (RS-2024-00333346 to J.-J.S.), the InnoCORE program of the Ministry of Science and ICT (GIST InnoCORE KH0860 to J.-J.S.) and National Institutes of Health (R01GM100108 to S.S.). K.-S.H. is supported by Brain Korea 21 program. The cryo-EM data were collected at KAIST Analysis Center Research Advancement (KARA). The computing resource was supported by GSDC and KREONET.

## Author Contributions

K.S., J.K., S.S., and J.-J.S. conceptualized the idea and K.S. and J.K. performed the experiments. S.S. and J.-J.S. supervised the experiments. All authors reviewed the data and wrote the manuscript.

